# AgRP neurons mediate activity-dependent development of oxytocin connectivity and autonomic regulation

**DOI:** 10.1101/2024.06.02.592838

**Authors:** Jessica E. Biddinger, Amanda E.T. Elson, Payam A. Fathi, Serena R. Sweet, Katsuhiko Nishimori, Julio E. Ayala, Richard B. Simerly

## Abstract

During postnatal life, the adipocyte-derived hormone leptin is required for proper targeting of neural inputs to the paraventricular nucleus of the hypothalamus (PVH) and impacts the activity of neurons containing agouti-related peptide (AgRP) in the arcuate nucleus of the hypothalamus. Activity-dependent developmental mechanisms are known to play a defining role during postnatal organization of neural circuits, but whether leptin-mediated postnatal neuronal activity specifies neural projections to the PVH or impacts downstream connectivity is largely unexplored. Here, we blocked neuronal activity of AgRP neurons during a discrete postnatal period and evaluated development of AgRP inputs to defined regions in the PVH, as well as descending projections from PVH oxytocin neurons to the dorsal vagal complex (DVC) and assessed their dependence on leptin or postnatal AgRP neuronal activity. In leptin-deficient mice, AgRP inputs to PVH neurons were significantly reduced, as well as oxytocin-specific neuronal targeting by AgRP. Moreover, downstream oxytocin projections from the PVH to the DVC were also impaired, despite the lack of leptin receptors found on PVH oxytocin neurons. Blocking AgRP neuron activity specifically during early postnatal life reduced the density of AgRP inputs to the PVH, as well as the density of projections from PVH oxytocin neurons to the DVC, and these innervation deficits were associated with dysregulated autonomic function. These findings suggest that postnatal targeting of descending PVH oxytocin projections to the DVC requires leptin-mediated AgRP neuronal activity, and represents a novel activity-dependent mechanism for hypothalamic specification of metabolic circuitry, with consequences for autonomic regulation.

**Significance statement:** Hypothalamic neural circuits maintain homeostasis by coordinating endocrine signals with autonomic responses and behavioral outputs to ensure that physiological responses remain in tune with environmental demands. The paraventricular nucleus of the hypothalamus (PVH) plays a central role in metabolic regulation, and the architecture of its neural inputs and axonal projections is a defining feature of how it receives and conveys neuroendocrine information. In adults, leptin regulates multiple aspects of metabolic physiology, but it also functions during development to direct formation of circuits controlling homeostatic functions. Here we demonstrate that leptin acts to specify the input-output architecture of PVH circuits through an activity-dependent, transsynaptic mechanism, which represents a novel means of sculpting neuroendocrine circuitry, with lasting effects on how the brain controls energy balance.

## Introduction

Early postnatal life is a period of dynamic brain growth and development, as young animals adapt to rapidly changing homeostatic needs and metabolic challenges. Accordingly, neural systems develop in response to metabolic and neuroendocrine sensory inputs that reflect the changing environment and nutritional conditions. In addition to its role of coordinating energy intake and expenditure in response to changes in nutritional status during adulthood, the adipocyte-derived hormone leptin functions as a developmental factor impacting the organization of neural pathways in the brain that control energy balance (1–5). Leptin signaling is required for postnatal axon outgrowth from the arcuate nucleus of the hypothalamus (ARH) to targets in the forebrain (6), including the paraventricular hypothalamic nucleus (PVH). Leptin also specifies distinct cellular targeting patterns to functionally defined compartments of the PVH, as well as cell type-specific targeting of ascending viscerosensory afferents from the nucleus of the solitary tract (NTS) to subpopulations of neurons in the PVH (7, 8). These neurotrophic actions of leptin are distinct from its regulatory role in adults and appear to be restricted to a discrete postnatal critical period that ends during the fourth postnatal week (9).

In addition to secreted growth factors, axon guidance cues, and circulating molecules such as leptin, neuronal activity can also profoundly impact the development of neural circuits. This phenomenon has been studied extensively in the visual system, where early patterns of neuronal activity exert significant lasting effects on ocular dominance (10); however, it is not known whether postnatal neuronal activity plays a similar role in specifying the architecture of hypothalamic circuitry. Although it is well established that development of AgRP inputs to preautonomic components of the PVH are dependent on postnatal environmental nutritional conditions (11, 12), development of projections from the PVH to the dorsal vagal complex (DVC) in the hindbrain has not been explored in this context (13, 14).

Oxytocin neurons in caudal regions of the PVH coordinate various aspects of ingestive behavior and autonomic regulation (15–23). Oxytocin neural circuits mature relatively late in development, as mature oxytocin peptide is not detected until after birth, and axon outgrowth progressively increases during the first few weeks of postnatal life (24–26). Descending oxytocin projections that arise from caudal PVH neurons and target the DVC are negligible at postnatal day 0 (P0), but markedly increase in density postnatally, reaching adult-like levels by weaning (27). Moreover, the postnatal organization of oxytocin neural circuits are broadly impacted by a variety of factors during development that can contribute to several neurodevelopment disorders, including Prader-Willi syndrome, Fragile X syndrome, autism spectrum disorders, and others (28–32).

Here, we assessed whether postnatal activity of AgRP neurons impacts innervation of oxytocin neurons in the PVH, and if AgRP activity or leptin are required for development of descending PVH oxytocin projections to the DVC. The results suggest that not only is leptin required for specifying oxytocin projections from the PVH to the DVC, but this developmental action of leptin on oxytocin neurons is receptor-independent and requires AgRP neuronal activity, suggesting that the input-output architecture of oxytocin neurons is developmentally specified by leptin through an activity-dependent transsynaptic mechanism.

## Results

### Activity-dependent development of AgRP projections to the PVH

Previous studies have established that axonal projections from AgRP neurons reach their target destinations during the first two weeks of postnatal life and that leptin signaling during a restricted postnatal sensitive period is required to achieve the full adult distribution (6, 33). Because neural activity is also known to regulate the establishment and refinement of neural circuits, we used a chemogenetic approach to inhibit AgRP neuronal activity during the critical period for development of AgRP projections to the PVH. Inhibitory DREADD (designer receptors exclusively activated by designer drugs) hM4Di receptors were selectively targeted to AgRP neurons (AgRP-Cre-hM4Di mice). Daily postnatal injections of clozapine N-oxide (CNO; 1.0 mg/kg i.p.) were administered from P4 to P14 (Fig. 1A). Induction of the inhibitory hM4Di DREADD signaling pathway resulted in labeling of AgRP-Cre-hM4Di neurons with the fluorescent reporter mCitrine by P10 (Fig. 1B). Most of the mCitrine-labeled AgRP-Cre-hM4Di-expressing neurons in the arcuate nucleus were also responsive to leptin at this age, consistent with previous reports showing AgRP neurons are directly responsive to leptin during postnatal life. Postnatal CNO administration reduced the number of leptin-induced cFos-immunoreactive nuclei in the ARH of AgRP-Cre-hM4Di mice by 67%, confirming effectiveness of CNO-mediated neuronal inhibition of AgRP neurons in postnatal mice (Fig. 1B). Notably, DREADD-mediated AgRP neuronal inhibition significantly reduced the density of AgRP-immunolabeled fibers in the PVH of adult AgRP-Cre-hM4Di mice, compared with that of saline-injected control mice (Fig. 1C-D), suggesting that development of these projections is dependent on the activity of AgRP neurons during postnatal life.

**Figure 1.**
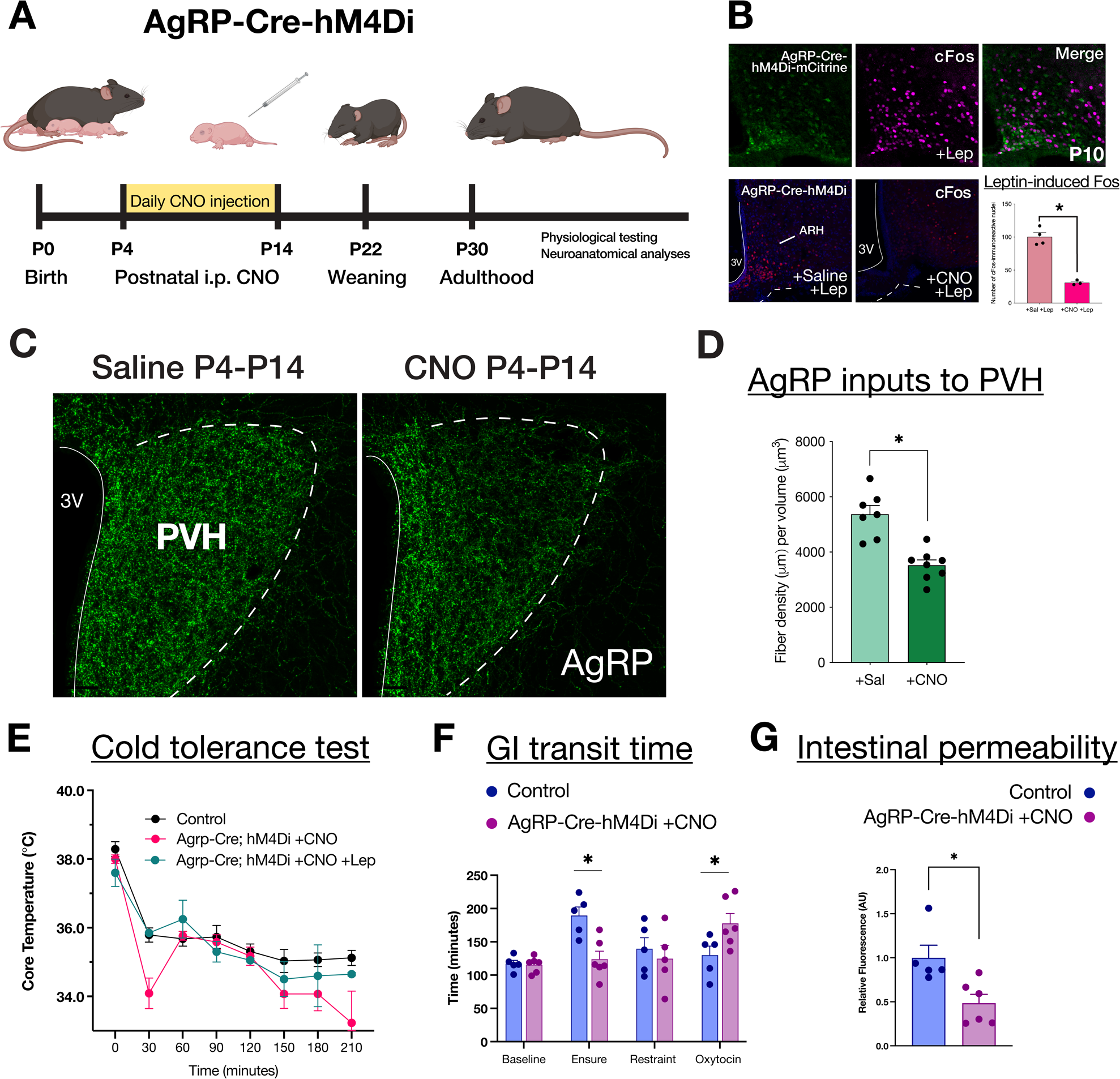
Postnatal activity of AgRP neurons specifies axon outgrowth and regulates autonomic output. Schematic of experimental paradigm. Mice received daily CNO (1.0 mg/kg i.p.) injections from P4-P14, and histology and autonomic function experiments were conducted in adulthood unless denoted otherwise (A). Confocal images illustrating ARH AgRP-Cre-hM4Di-mCitrine expression and localization of leptin-induced cFos at P10 (B). Postnatal DREADD-mediated AgRP neuronal inhibition results in significantly decreased AgRP fiber density in the PVH at P30, compared with control AgRP-Cre-hM4Di mice that received postnatal i.p. saline injections (C-D). Early postnatal AgRP neuron inhibition also impaired adult thermoregulation in response to a cold challenge (E), as well as dysregulated GI transit time in response to multiple GI metabolic stimuli (F) and altered intestinal permeability (G).

### Postnatal inhibition of AgRP neuronal activity is associated with autonomic dysfunction

Disrupted development of AgRP projections to presympathetic PVH neurons is associated with autonomic neuropathy (7). To test whether postnatal manipulation of AgRP neuronal activity impacts autonomic function, we measured physiological responses in adult AgRP-Cre-hM4Di mice that received postnatal injections of saline or CNO but remained untreated after P14 when exposed to a cold environment. In response to cold challenge, the ability of adult AgRP-Cre-hM4Di mice to maintain core body temperature was severely impaired in mice that received postnatal CNO treatment (Fig. 1E). All mice had similar core temperatures before cold exposure, and after 30 min at 4℃, the core temperature of all mice started to significantly decrease. However, the ability of AgRP-Cre-hM4Di +CNO animals to maintain their core body temperature was severely impaired; by 30 min their core temperatures decreased to half that of all other groups. As the duration of cold exposure increased, all mice began to lose the ability to regulate core body temperature, and after 180 minutes, AgRP-Cre-hM4Di +CNO animals were no longer able to maintain their core temperature. However, concomitant daily treatment with leptin and CNO from P4-P14, normalized core temperature of AgRP-Cre-hM4Di mice to that of controls, indicating that exogenous leptin administration during postnatal life can at least partially rescue activity-dependent defects and sympathetic responses.

The impact of silencing postnatal AgRP activity on gastrointestinal (GI) reflexes was tested by measuring GI transit time (GITT) in response to metabolic stimuli that are known to impact GI physiology. Oral gavage of Ensure led to longer GITT, but this response was blunted in AgRP-Cre-hM4Di mice that were treated postnatally with CNO. Furthermore, while acute oxytocin injection did not impact the GITT of adult mice in the fed state on a chow diet, GITT of AgRP-Cre-hM4Di mice treated with postnatal CNO significantly increased upon oxytocin administration (Fig. 1F). In addition, the gut epithelial wall of CNO-treated AgRP-Cre-hM4Di mice appeared to be less permeable to macronutrient content, as evidenced by a significant decrease in the concentration of FITC-labeled dextran in blood plasma after administration by oral gavage, indicating that the absorption of nutrients may be functionally impaired in adult mice that experienced reduced activity of AgRP neurons during postnatal life (Fig. 1G).

### Leptin is required for postnatal targeting of preautonomic PVH oxytocin neurons by AgRP neurons

Leptin promotes axonal outgrowth and cellular targeting of AgRP projections to functionally and cytoarchitectonically identified compartments of the PVH (7), but whether leptin specifies AgRP inputs to neurochemically-defined cell types is unknown. Oxytocin neurons located in parvocellular divisions of the caudal PVH send projections to medullary regions that have been identified as having functional significance for interoception. Therefore, we quantified AgRP-immunoreactive axons in close apposition to oxytocin neurons in distinct PVH compartments in *Lep^ob/ob^* mice and WT littermate controls. Consistent with previous results, AgRP inputs to the dorsal, lateral, and ventral medial parvocellular parts of the caudal PVH were significantly reduced in adult leptin-deficient animals (Fig. 2A, D, G). Moreover, by P30 the density of AgRP-immunoreactive terminals closely apposed to oxytocin neurons was significantly reduced in leptin-deficient mice compared with controls (Fig. 2C, F, I), indicating that leptin is required for cell-type specific targeting of AgRP inputs to oxytocin neurons in the PVH. This reduction in innervation density did not appear to be due to changes in the number of oxytocin neurons, as there was no difference in the number or volume of PVH oxytocin neurons in *Lep^ob/ob^* and WT mice (Fig. 2B, E, H).

**Figure 2.**
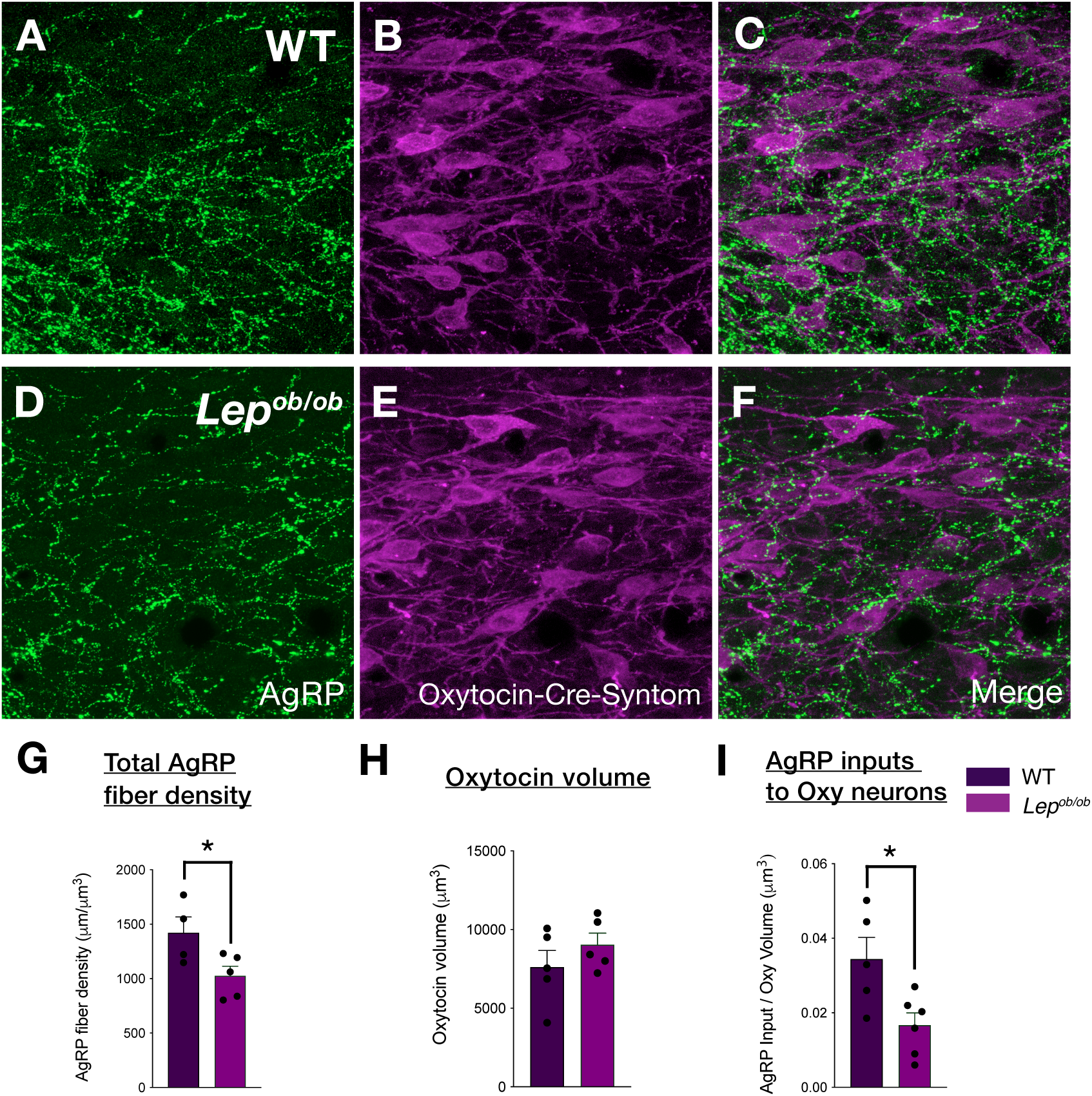
Leptin specifies targeting of AgRP projections to PVH oxytocin neurons. Confocal images depicting AgRP-immunoreactive axons (green) and Oxytocin-Cre-SynTom fluorescence (magenta) in Oxy-Cre-SynTom-WT (A-C) and leptin-deficient Oxy-Cre-SynTom-OB mice (D-F) in the PVH at P30 (A-B). Density of total AgRP-immunoreactive axons within the PVH was significantly decreased in leptin-deficient mice (A, D, G). AgRP-immunoreactive axons targeting Oxytocin-Cre neurons were significantly decreased in leptin-deficient mice compared with WT controls (C, E, I). No differences were observed in oxytocin volume (B, E, H).

Because AgRP innervation of presympathetic PVH neurons, including parvocellular oxytocin neurons, is especially sensitive to postnatal leptin, we used Cre-dependent, AAV-mediated axonal tracing in adult Oxytocin-Cre mice to map the projections of these neurons in the dorsomedial medulla (Fig. 3F). Viral transduction and location of AAV injection was confirmed by colocalization of tdTomato fluorescence and oxytocin immunolabeling (Fig. 3F-I) at the injection site. In these mice, we observed tdTomato-labeled axons in the DVC (Fig. 3J-L), which is comprised of the NTS, the dorsal motor nucleus of the vagus nerve (DMX), and the area postrema (AP). The tdTomato-labeled Oxytocin-Cre axonal inputs were not uniformly distributed within the DVC and appeared to be cellularly targeted within specific regional domains of the DVC and their subnuclei. The medial, gelatinous, and commissural subnuclei of the NTS at the level of the AP contained the highest density of tdTomato-labeled oxytocin axons (Fig. 3J-L), regions which receive input from vagal afferents, and are associated with viscerosensory transmission, especially gastric functions (27). At the level of the hindbrain with an open 4^th^ ventricle, just anterior to the rostral appearance of the AP, the dorsomedial NTS also received substantial oxytocin innervation (Fig. 3J). The NTS was generally more densely innervated by PVH oxytocin axons than the DMX, while the AP and underlying subpostrema area exhibited minimal axonal labeling (Fig. 3K, L). At rostral levels of the DMX, a considerable number of inputs were found along the dorsolateral edges of the medial gastric columns and the lateral celiac columns at (Fig. 3J-K). At mid-AP levels, (Fig. 3K), within the most lateral part of the medial NTS, along the dorsolateral edge of the underlying DMX, a very dense oxytocin terminal field was observed, near the region where the rostral A2 noradrenergic neurons and the caudal C3 adrenergic neurons are located. A few scattered axons were seen in NTS subnuclei lateral to the solitary tract (ts).

**Figure 3.**
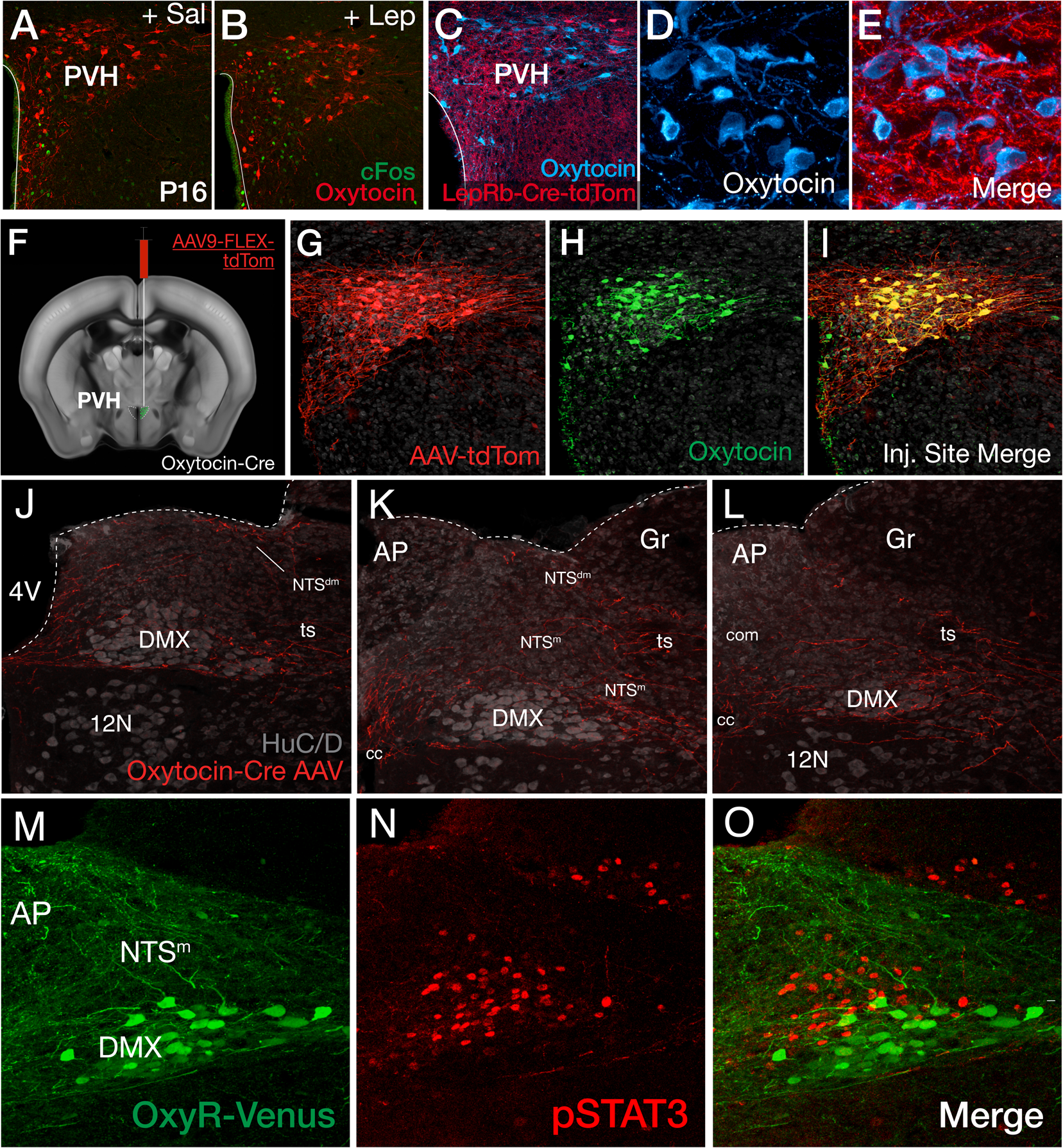
Leptin signaling in oxytocin and oxytocin receptor-expressing neurons in the PVH and DVC. Confocal images of oxytocin (red) and cFos (green) immunolabeling in the PVH of mice injected with i.p. saline or leptin at P16 (A, B). Representative confocal images of oxytocin-immunolabeled neurons (blue) and LepRb-Cre-tdTom fluorescence (red) in caudal regions of the PVH at P18, taken with 20x (C) and high-magnification 63x objectives, respectively (D, E). Stereotaxic injection of Cre-dependent AAV-tdTomato virus into the PVH of adult Oxytocin-Cre mice labels projections of neurons expressing Oxytocin-Cre (F). Confocal images depicting representative injection site of AAV-tdTomato virus (red) into Oxytocin-Cre-expressing neurons and oxytocin immunolabeled neurons (green) in the caudal PVH (G-I). TdTomato-labeled Oxytocin-Cre axonal inputs (red) are shown at three levels of the dorsomedial medulla after AAV-mediated recombination. Neurons (gray) are immunolabeled with the pan-neuronal marker HuC/D (J-L). Confocal image of OxyR-expressing neurons in the DVC at P30, visualized by OxyR-Venus fluorescence (M). Representative image of pSTAT3-immunoreactivity (red) in the DVC of OxyR-Venus mice injected i.p. with leptin (N). Merged image showing pSTAT3-immunoreactive nuclei in response to i.p. leptin injection and DVC OxyR-Venus labeled neurons (O). Abbreviations: PVH; paraventricular nucleus of the hypothalamus; DVC, dorsal vagal complex; AP, area postrema; DMX, dorsal motor nucleus of the vagus nerve; Gr, Gracile nucleus; NTSm, medial subnucleus of the solitary tract; NTSdm, dorsomedial subnucleus of the solitary tract; 3V, third ventricle; cc, central canal; ts, solitary tract.

### Leptin and postnatal development of descending PVH oxytocin projections to the dorsal vagal complex

To determine if PVH oxytocin neurons are sensitive to the postnatal actions of leptin, mice received acute injection of either saline or leptin (10 mg/kg i.p.) at P16 and subsequently perfused for immunohistochemical localization of cFos. Leptin treatment resulted in a moderate number of cFos-immunoreactive nuclei in multiple PVH subcompartments, primarily the dorsal, ventral, and medial parvicellular regions (Fig. 3A-B). These functionally-defined compartments are located in the caudal PVH and project to the DVC, where they are known to regulate GI functions (34–37). However, although leptin-induced cFos was observed in the PVH at P16, it was virtually absent within oxytocin neurons (Fig. 3A-B). The density of oxytocin-immunolabeled neurons that also co-express LepRb-targeted tdTomato was also low at P16, suggesting that leptin receptors are not broadly expressed on PVH oxytocin neurons during postnatal life. However, PVH oxytocin neurons appear to be densely innervated by axons containing LepRb-Cre-tdTomato fluorescence, presumably derived from neuronal populations that provide direct inputs to PVH oxytocin neurons (Fig. 3C-E). Although PVH oxytocin neurons do not appear to express LepRb, and postnatal leptin does not directly stimulate these neurons, development of descending oxytocin projections may take place through a target-dependent mechanism. To explore the possibility that oxytocin receptor-expressing neurons in the DVC may be directly activated by leptin during postnatal development, mice with Oxytocin Receptor-Venus fluorescence were assessed for leptin-induced phospho-STAT3 (pSTAT3)-immunoreactivity. Interestingly, robust induction of pSTAT3-immunoreactivity was localized to the medial and commissural NTS subnuclei, as well as the adjacent gracile nucleus, while pSTAT3 was absent in oxytocin receptor neurons, suggesting that, like PVH oxytocin neurons, oxytocin receptor-expressing DVC neurons are not directly responsive to leptin (Fig. 3M-O).

### Activity-dependent development of oxytocin inputs to the DVC

To compare development of oxytocin projections to the DVC in WT and *Lep^ob/ob^* mice, oxytocin axons were visualized by genetically targeting the synaptophysin-tdTomato axonal reporter to Oxytocin-Cre expressing neurons of WT (Oxytocin-Cre-SynTom-WT) and leptin-deficient (Oxytocin-Cre-SynTom-OB) mice at multiple ages (Fig. 4A-L). Oxytocin-Cre-SynTom labeled projections to the DVC were markedly reduced in *Lep^ob/ob^* mice by P16 (Fig. 4B, F, J), and remained low into adult life at P30 (Fig. 4K) and P60 (Fig. 4L). No changes in the number of Oxytocin-Cre-SynTom labeled neurons in the PVH were detected at P16 or P30 (Fig. 4M-Q). The leptin-dependent innervation of PVH oxytocin neurons by AgRP neurons, which are highly responsive to leptin during postnatal life, may provide a transsynaptic activity impacting development of descending oxytocin projections to the DVC. To test this, we assessed the PVH oxytocin projections to the DVC, using DREADD-mediated inhibition to block the activity of AgRP neurons during early postnatal life. Daily postnatal CNO treatments to AgRP-Cre-hM4Di mice resulted in a significant reduction in the density of oxytocin projections to the DVC in adult mice (Fig. 4R-T). Thus, in contrast to the direct action of leptin on targeting AgRP axonal projections to the PVH, these findings suggest that postnatal specification of descending PVH oxytocin projections to the DVC is dependent on an indirect mechanism that requires AgRP neuronal activity.

**Figure 4.**
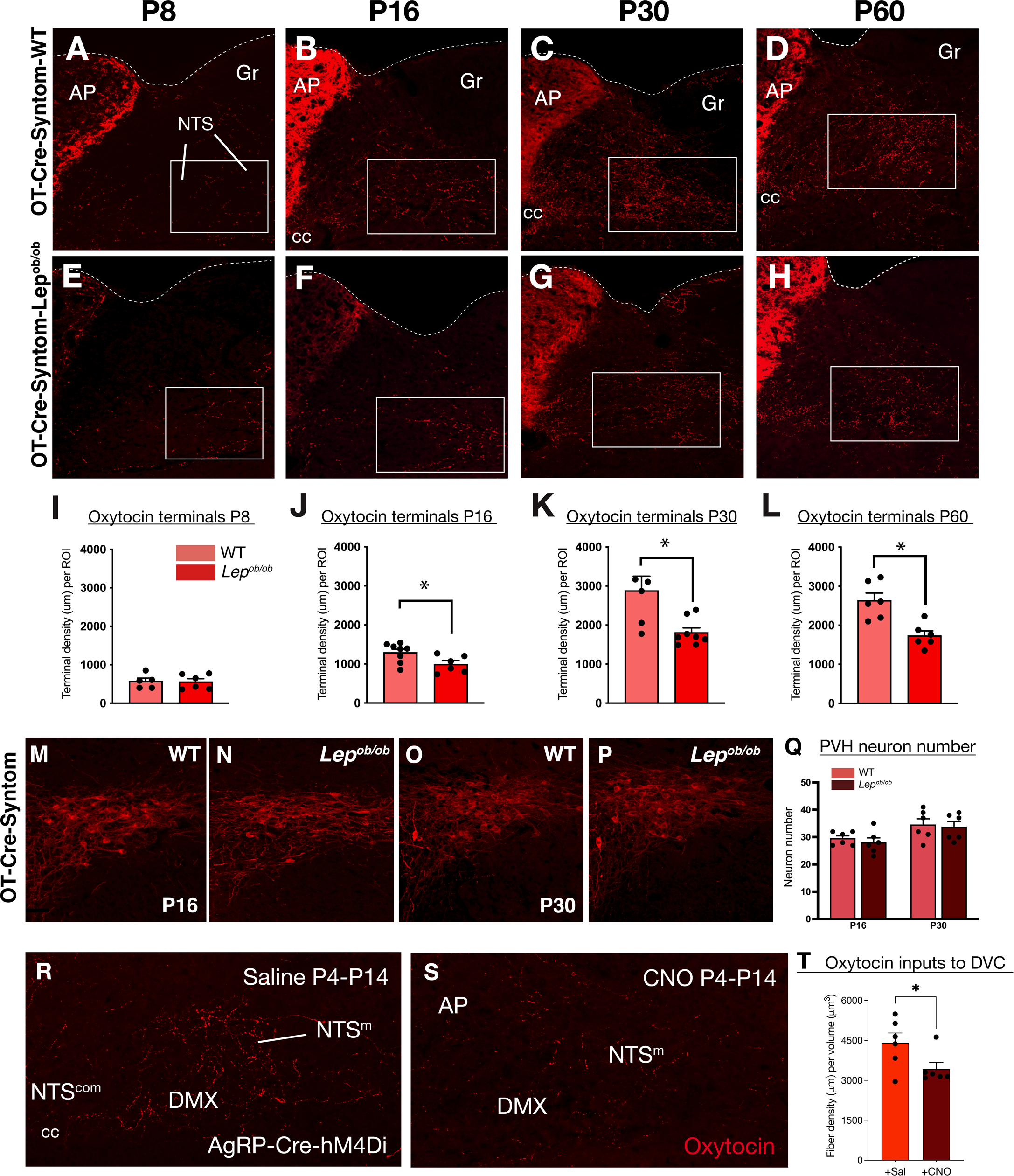
Altered postnatal neuronal signaling impairs projections to the dorsal vagal complex. Oxytocin projections to the DVC were labeled by targeting fluorescent synaptophysin-tdTomato to Oxytocin-Cre axons (A-L) and neurons (M-Q). Confocal images of oxytocin-immunoreactive fibers at P30 in the DVC of AgRP-Cre-hM4Di mice that received injections of either saline or CNO i.p. (R-T) from P4-P14.

## Discussion

As a central regulator of energy balance, the PVH integrates multiple sources of hormonal, viscerosensory and extrasensory information to effectively coordinate metabolic state with feeding behavior and autonomic regulation. Key among these signals is leptin, which informs the brain about the status of energy stores in peripheral tissues (1, 38). Subpopulations of PVH neurons impact multiple aspects of autonomic regulation and feeding behavior through direct projections to regions such as the NTS, parabrachial nucleus and lateral hypothalamus (34, 35, 39–42). Thus, changes in the input-output architecture of the PVH will likely alter how it functions to integrate these diverse signals and regulate downstream effectors of neuroendocrine and autonomic function.

Targeting of AgRP inputs to preautonomic PVH neurons, but not to neuroendocrine neurons in the PVH, can be sustainably rescued by treatment of *Lep^ob/ob^* mice with exogenous leptin during the first two weeks of postnatal life (6, 7). During this same period, targeting of GLP-1 afferents from the NTS to specific subpopulations of PVH neurons are likely mediated by cell autonomous leptin receptor signaling (8, 43). For example, in adult *Lep^ob/ob^* mice there appears to be a reduction in AgRP inputs to oxytocin neurons in the PVH, but no change in ascending viscerosensory inputs from NTS PPG neurons to PVH oxytocin neurons, (8) suggesting that the developmental actions of leptin can permanently specify the balance in sensory information normally conveyed to oxytocin neurons by at least these two afferent pathways.

Data from our AAV tracing experiments confirm that descending projections from PVH oxytocin neurons provide direct inputs to distinct regions of the DVC, where they influence several aspects of autonomic function by modulating the impact of incoming visceral sensory information (16–18, 21, 27, 36, 40, 41, 44–53). These descending oxytocin projections appear to be established during the postnatal period (27), when animals begin to explore solid food in preparation for independent ingestion (54), and when the postnatal leptin surge functions to impact the organization of afferents to the PVH (55). Our analysis of oxytocin projections to the DVC in *Lep^ob/ob^* mice suggest that leptin is required for postnatal targeting of these projections, without affecting the number of PVH oxytocin neurons. However, in contrast to the receptor-mediated developmental actions of leptin on AgRP and GLP-1 afferents to the PVH, postnatal development of descending oxytocin projections to the DVC appears to be independent of LepRb-mediated signaling in oxytocin neurons, because these cells do not display LepRb expression or STAT3 activation in response to postnatal leptin treatment. This observation suggests an indirect mechanism of leptin action on neural development, perhaps through activity-dependent activation of axon targeting. Thus, leptin signaling appears to function indirectly during postnatal life to alter development of PVH oxytocin neurons, in contrast to its regulatory actions documented previously in adult rodents (16, 22, 56–58).

Neuronal activity has not been shown previously to impact development of hypothalamic circuits, however, activity-dependent axonal targeting has been well documented in other neural systems. Studies in the visual system of multiple species have demonstrated that neural activity, either spontaneous or patterned, is required for normal development of central projections (10, 59, 60). Similarly, patterns of innervation in cerebellum and cerebral cortex are also dependent on neural activity (61–63). Based on our observations, development of AgRP inputs to the PVH and downstream targeting of oxytocin projections to the DVC appear to depend on the postnatal activity of AgRP neurons. Chemogenetic silencing of AgRP neuronal activity during the developmental critical period for innervation of PVH neurons caused a lasting reduction in the density of AgRP inputs in adult mice. Peripubertal silencing of AgRP neuronal activity using chemogenetics impacts the structure and function of precortical cells, as well as food intake (64). Together with previous results in *Lep^ob/ob^* mice (6), our findings suggest that perturbations in leptin, or other signals that alter firing of AgRP neurons during postnatal development, may lead to permanent changes in innervation patterns in the PVH.

The anatomical changes in descending projections from PVH oxytocin neurons to DVC targets correspond to disruptions in cold tolerance and GI motility, thereby raising the possibility that postnatal AgRP activity may exert a lasting effect on autonomic function. Consistent with this interpretation, PVH oxytocin neurons have been implicated in control of adaptive thermogenesis and energy expenditure, both of which are impacted by leptin signaling (65–68). Moreover, neurons in the PVH innervate preganglionic neurons to influence outflow to the stomach, and oxytocin alters vagal efferents and impacts gastric motility (34, 53, 69–71).

Consistent with a modulatory role for oxytocin in gastric function (72), our data suggest that a decrease in oxytocin projections to the DVC is associated with longer GI transit time. Moreover, the structural dependency between AgRP activity and PVH oxytocin neurons also suggests a hypothalamic to brainstem pathway in which PVH oxytocin may function as a downstream mediator of hypothalamic leptin to regulate meal-related satiety signals and other GI functions, despite directly lacking leptin receptors.

Previous ex-vivo electrophysiological studies determined that leptin excites AgRP neurons during postnatal life, in contrast to its hyperpolarizing action in mature mice after they transition to independent feeding (73). Thus, the ability of leptin to activate AgRP neurons when their projections are extending into the PVH during postnatal development, may function as a temporally limited developmental signal analogous to retinal activity or experience during the critical period for development of the visual system. Moreover, this activity-dependent developmental action of leptin may extend to downstream targets involved in autonomic regulation. Impairment of oxytocin projections to the DVC caused by silencing AgRP neuron activity during the postnatal period suggests that activity of AgRP neurons may transynaptically impact development of descending projections from oxytocin neurons through an activity-dependent mechanism that is independent of cell autonomous LepRb signaling. Thus, in addition to direct receptor-mediated actions of leptin on development of AgRP projections to the PVH, regulatory factors that alter the activity of AgRP neurons during this postnatal critical period may exert a lasting impact on targeting of AgRP axons in the PVH, as well as on development of downstream projections of cells innervated by AgRP neurons. Although determining how leptin calibrates topographic maps of AgRP innervation to functionally distinct components of the PVH will require further experimentation, the findings reported here support the notion that activity-dependent patterning of AgRP inputs to the PVH, and subsequent development of oxytocin projections from the PVH to the DVC, represent a novel role for leptin in specifying development of neural circuits involved in autonomic regulation and food intake.

## Materials and Methods

Please see SI Appendix for detailed methodological information.

### Animals

Mice were housed at 22°C on a 12 hr light-dark cycle and provided ad libitum access to a standard chow diet (PicoLab Rodent Diet 20 #5053) and water. Mice were weaned at P22 and maintained in their home cages with mixed genotype littermates until they were used for experiments. All animal care and experimental procedures were approved by the Institutional Care and Use Committee of Vanderbilt University and the Saban Research Institute at Children’s Hospital Los Angeles. All procedures were performed in accordance with the guidelines of the National Institutes of Health.

### Experimental paradigms and analyses

Data are presented as group mean values ± SEM, as well as individual data points. Statistical analyses were performed using GraphPad Prism software (Version 9.5). Two-way repeated measures analysis of variance (ANOVA) with multiple comparisons was used to compare groups in the cold challenge assay. One-way ANOVA followed by a pairwise post-hoc test was used to test for comparisons between three groups. Student’s t-test was used to compare data within two groups, using paired and unpaired tests where appropriate. Differences between groups were considered statistically significant at *p*<0.05.

## Acknowledgements

This work was supported by National Institute of Health grant DK106476 to R.B.S. We thank all members of the Simerly Lab for helpful discussion and input on the manuscript, and especially Dollada Srisai for providing technological support. We would also like to thank Dong Zhou and Frohar Mirzai for animal care, and Nicholas Thomas-Low for excellent histochemical support. We thank Larry Young at Emory University for providing breeders to establish a local colony of Oxytocin-Receptor-Venus mice, and the Mouse Metabolic Phenotyping Core (MMPC) at Vanderbilt University.

## Competing Interest Statement

No competing interests

## Classifications

Major category: Biological Sciences; Minor category: Neuroscience

---

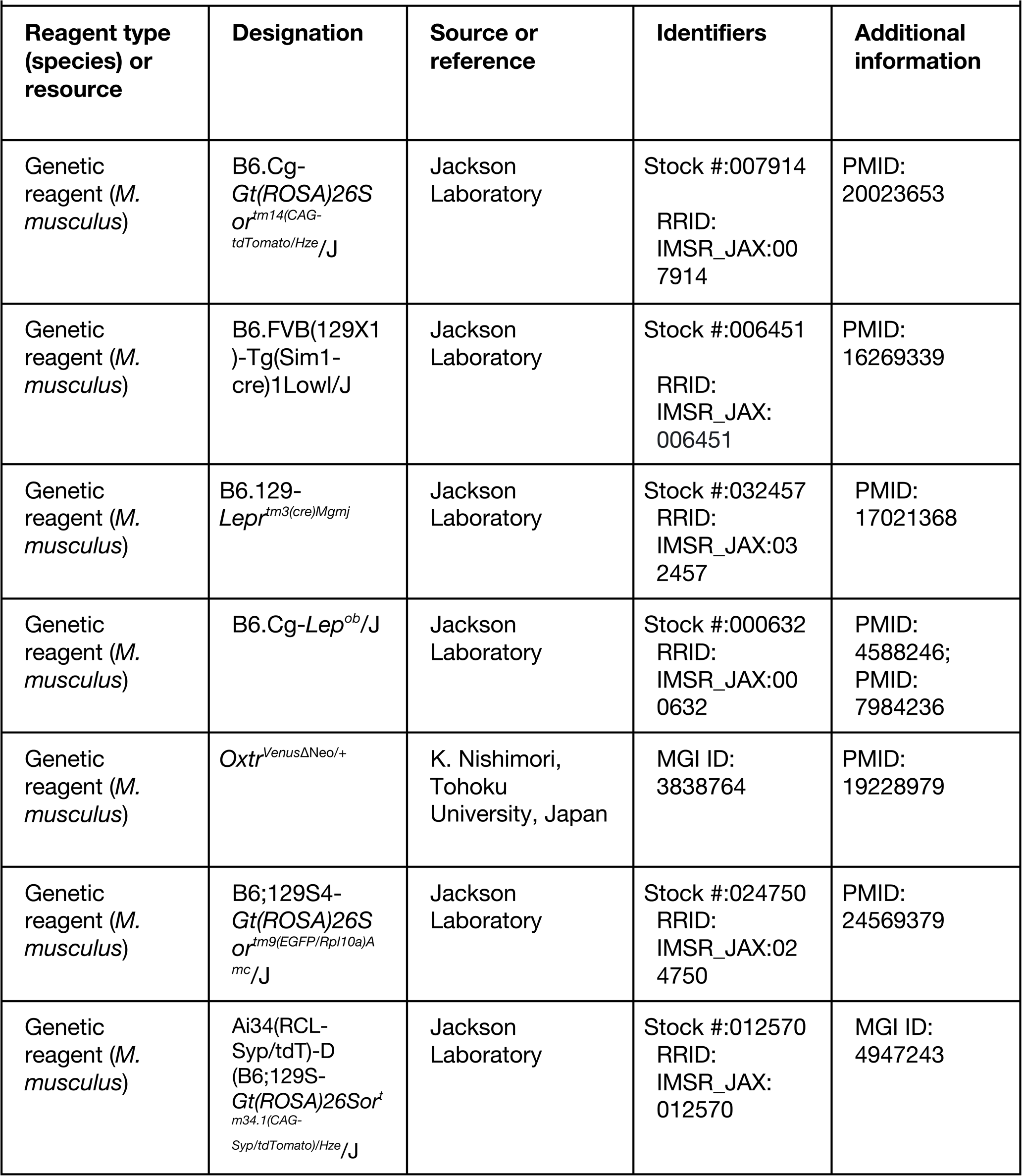
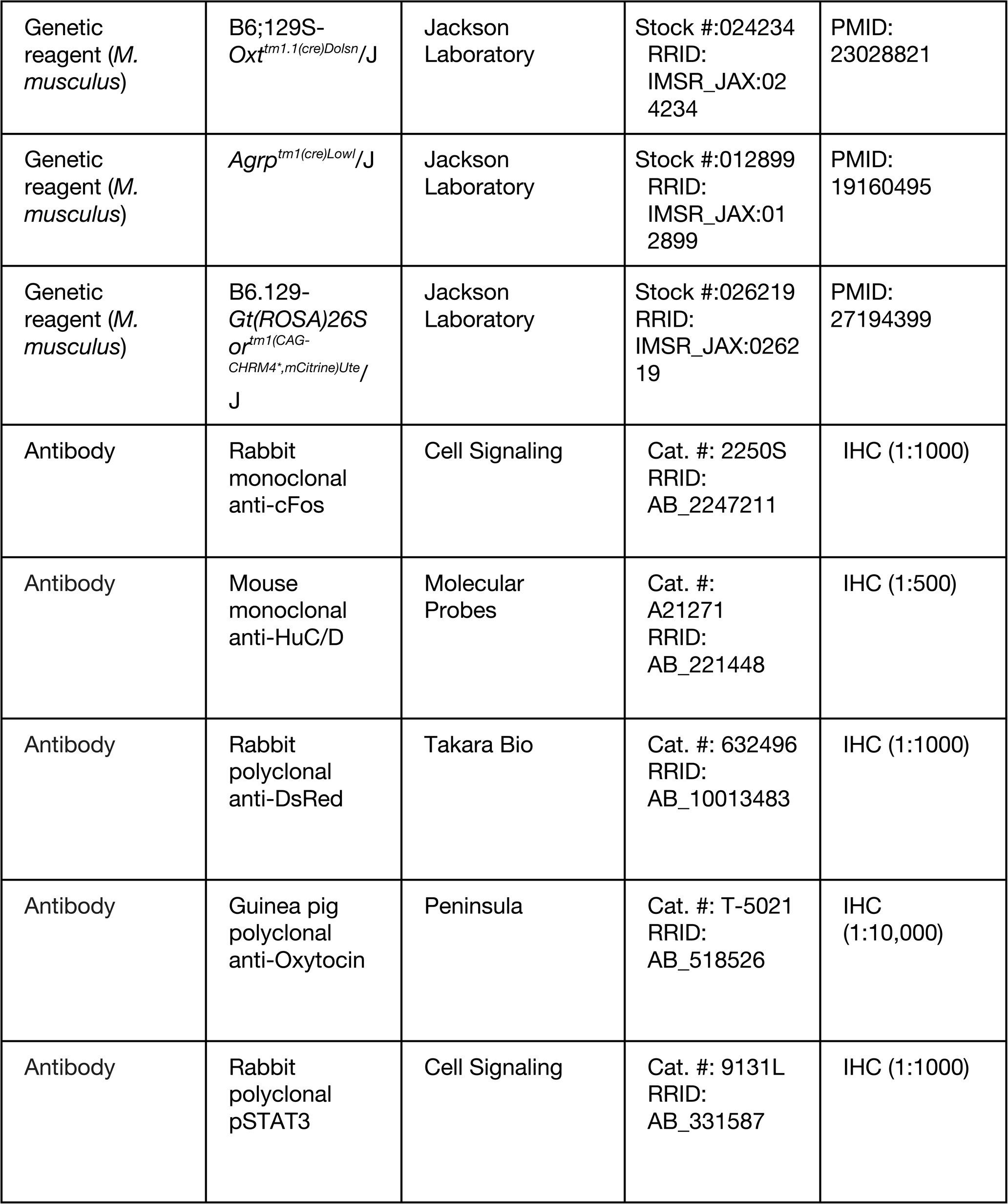
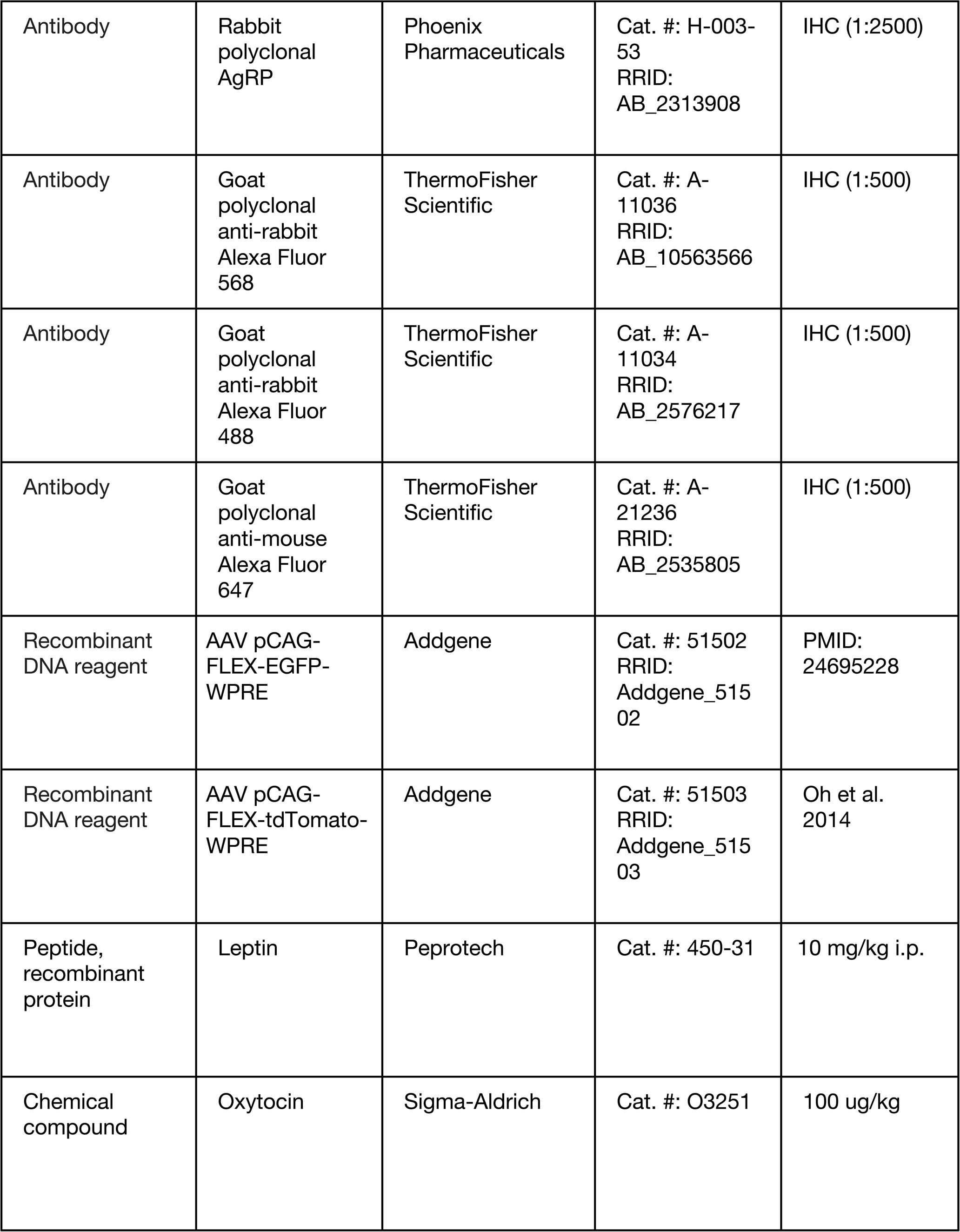
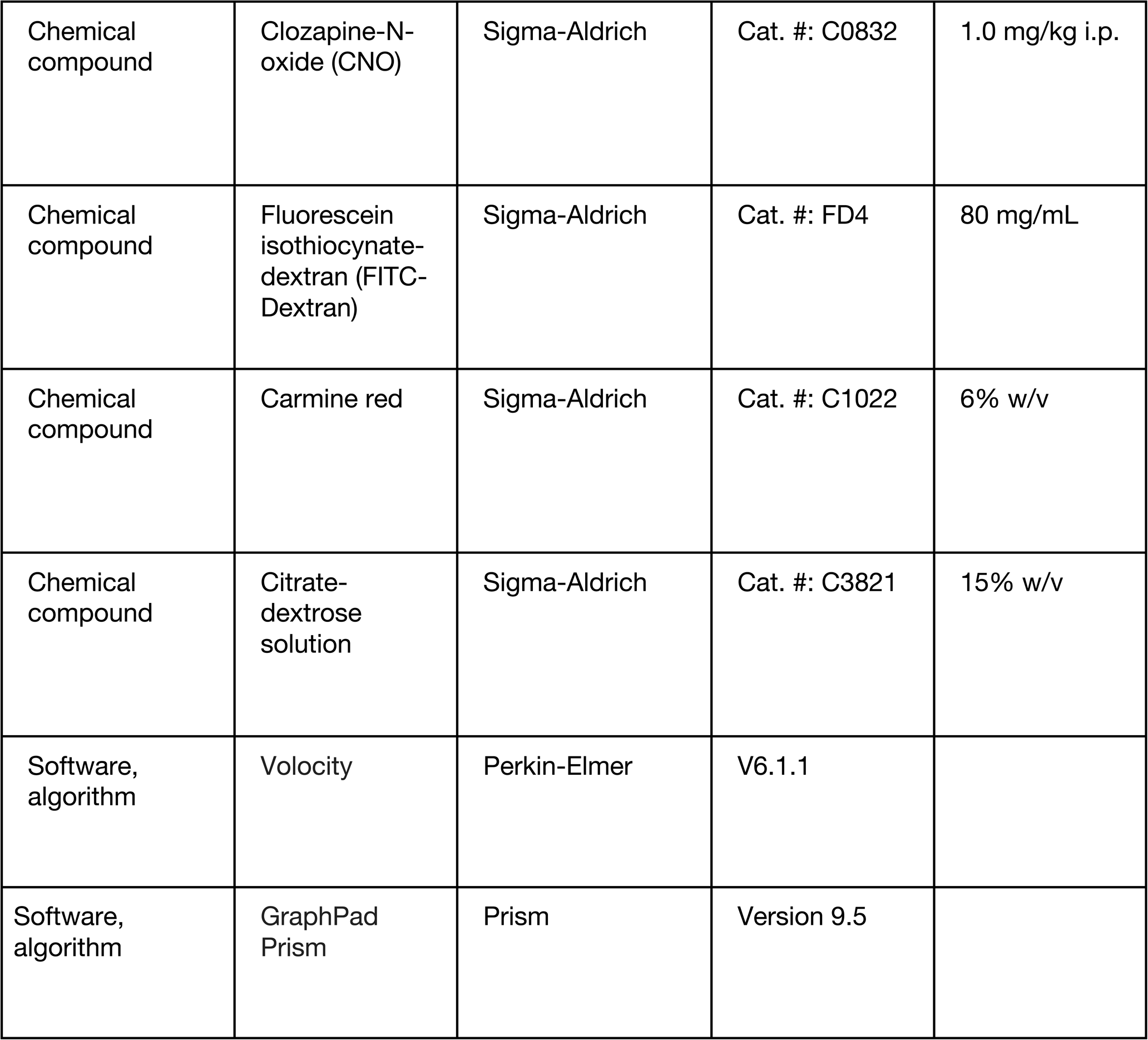
Key Resources Table

## Materials and Methods

### Animals

All animal care and experimental procedures were approved by the Institutional Care and Use Committee of Vanderbilt University and the Saban Research Institute at Children’s Hospital Los Angeles. All procedures were performed in accordance with the guidelines of the National Institutes of Health. Mice were housed at 22°C on a 12 hr light-dark cycle and provided ad libitum access to a standard chow diet (PicoLab Rodent Diet 20 #5053) and water. Mice were weaned at P22 and maintained with mixed genotype littermates until males were used for experiments.

Mouse lines used in this study: transgenic Sim1-Cre mice (B6.FVB(129X1)-Tg(Sim1-cre)1Lowl/J; stock number: 006451) were crossed with EGFP-L10a mice (B6;129S4-*Gt(ROSA)26Sor^tm9(EGFP/Rpl10a)Amc^*^/^J; stock number: 024750) to visualize constitutive expression of the L10a-EGFP fusion protein in Sim1-Cre-expressing neurons upon Cre-mediated excision of loxP-flanked transcriptional STOP fragments (Sim1Cre; EGFP-L10 mice) and used for stereotaxic surgery. Adult Oxytocin-IRES-Cre mice (B6;129S-*Oxt^tm1.1(cre)Dolsn^*^/^J; stock number: 024234) express Cre recombinase targeted to the endogenous oxytocin (*Oxt)* gene promoter and were used for stereotaxic surgery and histology. A separate cohort of Oxytocin-IRES-Cre mice were bred with mice heterozygous for the spontaneous obese (*ob*) mutation of the leptin-encoding gene (B6.Cg-*Lep^ob^*/J; stock number: 000632). The compound heterozygous offspring that result from this pairing (Oxytocin-Cre*; Lep^ob/+^* mice) were subsequently mated with double transgenic Ai34(RCL-Syp/tdT)-D (B6;129S-*Gt(ROSA)26Sor^tm34.1(CAG-Syp/tdTomato)/Hze^*/J; stock number: 012570); *Lep^ob/+^* mice in order to induce Cre-dependent expression of a Synaptophysin-tdTomato fusion protein (SynTom) targeted to neural elements that express Oxytocin-Cre in leptin-deficient mice (Oxytocin-Cre-SynTom-*Lep^ob/ob^*) and WT littermate controls (Oxytocin-Cre-SynTom-WT). Oxytocin receptor-Venus knock-in mice *(Oxtr^Venus^*^ΔNeo*/+*^) were used to visualize oxytocin receptor expression through Cre-mediated excision of a floxed PGK-Neo allele that drives expression of an enhanced YFP variant (Venus) under control of the endogenous regulatory sequence that encodes the oxytocin receptor gene (*Otxr*)(1). Leptin receptor-IRES-Cre knock-in mice (B6.129-*Lepr^tm3(cre)Mgmj^*; stock number: 032457) have Cre recombinase targeted to the long form of the leptin receptor gene-specific exon (LepRb-Cre mice) and were crossed with Cre-dependent Ai14(RCL-tdT)-D reporter mice (B6.Cg-*Gt(ROSA)26Sor^tm14(CAG-tdTomato/Hze^*/J; stock number: 007914) to visualize tdTomato3 fluorescent protein in neurons that express LepRb-Cre following Cre-mediated recombination (LepRb-Cre-tdTom mice). Agrp-IRES-Cre knock-in mice (*Agrp^tm1(cre)Lowl^*/J; stock number: 012899) express Cre recombinase in endogenous regulatory regions encoded by the *Agrp* gene. These animals were mated with R26-LSL-hM4Di-DREADD mice (B6.129-*Gt(ROSA)26Sor^tm1(CAG-CHRM4*,mCitrine)Ute^*/J; stock number 026219), that have targeted expression of a Cre-inducible genetic construct containing mutant hM4Di G protein-coupled receptor (GPCR) human M4 muscarinic (CHRM4) DREADD (designer receptor exclusively activated by designer drug) sequence, hemagglutinin epitope tag (HA), and yellow-green YFP variant mCitrine. Cre-mediated recombination of flanked loxP STOP sites results in transcription of HA-tagged hM4Di-mCitrine DREADD, targeted to neurons expressing AgRP-Cre (AgRP-Cre; hM4Di mice). Upon administration of DREADD ligand clozapine-N-oxide (CNO), the mutant hM4Di GPCR induces the canonical Gi-coupled inhibitory signaling pathway, in order to selectively inhibit AgRP neuronal activity. All mice were obtained from The Jackson Laboratory unless otherwise specified.

### Stereotaxic surgery

For viral tracing experiments, Cre-dependent AAV9-CAG-FLEX-WPRE viruses (Addgene) was injected into the PVH of Sim1-Cre; EGFP-L10 and Oxytocin-Cre mice. Stereotaxic coordinates used to target injections to the caudal PVH in adult and postnatal mice were determined from bregma using the Allen Reference Atlas of the C57BL/6J mouse brain (2), and are as follows: A/P, -0.90; M/L, ±0.2; D/V, - 5.0 from bregma (adults), and A/P, -0.50; M/L, ±0.1; D/V, -4.0 from bregma (P12 mice). Mice were first anesthetized with isoflurane (1.5-2.0%, 1L/minute in O2) and then placed into a stereotaxic frame (David Kopf Instruments). The skull was exposed through a small midline incision in the skin at the dorsal surface of the head, and the brain was accessed via a small hole at the surface of the skull. A fine-tipped glass micropipette filled with virus (inner diameter 20-30 µm) was slowly lowered into the brain and pressure-injected by using a PicroSpritzer system (General Valve Corporation; 40 psi, average duration: 4 msec). The tip of the glass micropipette was left in the brain for at least 5 minutes, and then slowly retracted. Vetbond was used to close the skin, and mice recovered from anesthesia on a heating pad. Mice were administered postoperative analgesia (Ketoprofen, 5 mg/kg) and monitored daily. Mice were allowed a minimum of two weeks for viral transduction and recovery from surgery. Accurate targeting of viral injections was determined by post-hoc histological assessment, and animals were only4 included in subsequent analyses if viral expression at the injection site was restricted to the PVH target region. Injection sites were also confirmed by co-immunolabeling with an antibody to oxytocin.

### Tissue Preparation

Mice were perfused and processed for histochemical experiments as previously described (3) unless otherwise specified. Briefly, mice were first anesthetized with tribromoethanol (TBE), then perfused transcardially with 0.9% saline, followed by ice-cold fixative (4% paraformaldehyde in 0.1M borate buffer, pH 9.5) for 20 min. Brains were carefully removed from the skull and postfixed in the same fixative for 4 hr at 4°C. After cryoprotection overnight in 20% sucrose, brains and spinal cords were frozen in powdered dry ice and stored at -80°C. A sliding microtome was used to collect 30 μm-thick coronal sections from adult mice (P30-P60) and free-floating tissue sections were stored in cryoprotectant solution at -20°C until further processing. As brains from postnatal mice (P8-P16) are smaller and more fragile, they were first embedded in OCT compound and a cryostat was used to collect 20 μm-thick coronal sections. The sections were mounted directly onto charged Superfrost Plus slides (Fisher Scientific), and after drying overnight at 4°C, slides were placed into cryoprotectant and stored at -20°C until further processing.

### Treatments and immunohistochemistry

To prepare tissue for immunohistochemical processing, tissue sections were removed from cryoprotectant and rinsed several times in 0.02M KPBS. Sections were incubated in a blocking buffer containing 2% normal goat or donkey serum and 0.3% Triton X-100 in 0.02M KPBS overnight at 4°C. Primary antibodies were diluted in the same blocking buffer described above, and used at the following concentrations: guinea pig anti-Oxytocin (1:10,000; Peninsula), rabbit anti-DsRed (1:1000; Takara Bio); mouse anti-HuC/D (1:500; Molecular Probes), rabbit anti-cFos (1:1000; Cell Signaling), rabbit anti-pSTAT3 (1:1000; Cell Signaling), chicken anti-GFP (1:10,000; Chemicon), rabbit anti-AgRP (1:2500; Phoenix Pharmaceuticals). Tissue sections were incubated in blocking buffer containing combinations of primary antibodies for 48 hr at 4°C. Primary antisera that were raised in the same species were not combined into solution, to ensure labeling specificity and avoid species cross-reactivity, and staining with heterologous combinations of primary and secondary antisera did not produce detectable staining. Sections were rinsed several times in 0.02M KPBS, and were then incubated in the appropriate species-specific fluorophore-conjugated Alexa5 Fluor secondary antibodies for 1 hr at RT. Tissue sections were again rinsed several times in 0.02M KPBS, and then coverslipped with No.1.5 Gold Seal cover glass (Electron Microscopy Services) and ProLong Gold antifade mounting medium (Invitrogen).

Staining for pSTAT3 or cFos nuclear immunoreactivity was performed as previously described (3) Briefly, age-matched groups of mice received intraperitoneal (i.p.) injections of recombinant mouse leptin (10 μg/g; Preprotech Inc) or 0.9% sterile NaCl on P10, P16, or P30. Mice were anesthetized with TBE 45 min after leptin injection for pSTAT3 immunolabeling, or 60 min later for cFos. All animals were perfused transcardially with 0.9% NaCl, followed by ice-cold fixative: 2% paraformaldehyde in 0.1M sodium phosphate buffer, pH 7.4 for 10 min (pSTAT3), or 4% paraformaldehyde in 0.1M borate buffer, pH 9.5 for 20 min if adults, and 10 min for pups (cFos). Brains were postfixed in the same fixative for 2 (pSTAT3) or 4 hr (cFos), and cryoprotected in 0.02M KPBS with 20% sucrose overnight. Each brain was then processed for either pSTAT3 or cFos immunohistochemistry. For pSTAT3 immunolabeling, tissue sections were first mounted onto Superfrost Plus slides, and then dried overnight at 4°C. The following day, sections were pretreated with 0.5% NaOH / H2O2, followed by incubation in 0.3% glycine, then 0.03% SDS, all made in 0.02M KPBS buffer. After these pretreatment steps, the mounted sections were incubated on slides in blocking buffer solution containing 4% normal goat serum, 0.4% Triton-X, and 1% BSA (fraction V) at RT for 20 min, followed by incubation in rabbit anti-pSTAT3 primary antibody (1:1,000; Cell Signaling) at 4°C for 48 hr. Sections were rinsed in 0.02M KPBS several times, and incubated for 2 hr at RT in the same blocking buffer described above, including goat anti-rabbit Alexa-Fluor conjugated secondary antibodies (1:500; Invitrogen), after which the sections were rinsed again with 0.02M KPBS, counter-stained with bis-benzamide (DAPI; 1:10,000; Invitrogen), and cover slipped as described above.

To inhibit AgRP neuronal activity in postnatal mice, AgRP-Cre-hM4Di mice and littermate controls received daily i.p. injections of either CNO (1.0 mg/kg; Sigma-Aldrich) or sterile saline from P4-P14. Mouse pups were weighed daily and remained in their home cage with their littermates and biological dam, with ad libitum access to standard rodent chow. Injections were administered in the home cage, at least 1 hr prior to lights out. Mouse pups were weaned onto the same chow diet at P22 and used for further6 experiments as adults. Upon completion of physiological testing in adulthood, mice were perfused and processed for immunohistochemistry as described above.

### Gastrointestinal transit time assay

Mice were singly housed, and were provided ad libitum access to a standard chow diet (PicoLab Rodent Diet 20 #5053) and water prior to the start of each experiment unless otherwise noted and returned to their respective home cages following the conclusion of each experiment. Whole gastrointestinal transit times were measured using a solution comprised of 6% Carmine Red (w/v) and 0.5% methylcellulose diluted in sterile saline. Mice received 250 uL of Carmine Red solution via oral gavage and the baseline gastrointestinal transit time was determined by collecting stools every 10-15 minutes until the first red colored stool was observed. We next examined the effects of nutritive preload oral gavage (0.5 mL Ensure per mouse) on gastrointestinal transit time in conditions with ad-lib access to food and following an 18-hour fast, and intraperitoneal oxytocin (100 ug/kg) administration on gastrointestinal transit times.

### Gastrointestinal permeability

To test whether postnatal AgRP neuronal activity functionally impacts GI physiology, we adapted a previously validated in vivo gut permeability assay (5) The absorption of fluorescein isothiocyanate (FITC)-labeled dextran was evaluated by measuring its concentration in blood plasma after administration by oral gavage. Mice were singly housed and fasted for four hours prior to oral gavage of .4 kDa FITC-dextran (80 mg/mL in PBS pH 7.4, 150 uL per mouse). Tail blood was collected just prior to and four hours after FITC-dextran administration. Plasma was recovered by mixing samples with Citrate-dextrose solution (15% v/v) and centrifuging at 5,000 rpm for 10 minutes. Plasma samples were diluted in 1X PBS (1:10 in duplicate) and relative fluorescein concentration was determined using a SpectraMax M4 microplate reader (excitation 485 nm, emission 530 nm). Mice were allowed to recover for at least 7 days prior to additional experiments.

### Cold tolerance test

Mice were placed in individual cages at 4℃ with ad-lib access to food and water. Core body temperature was recorded at time 0 (the onset of cold exposure) and then every 30 min of cold exposure, for a total of 210 min, using a rodent rectal probe (Physitemp).

### Image acquisition and analysis

First, brain tissue sections containing the hypothalamic arcuate and PVH nuclei, and caudal hindbrain sections that include the dorsomedial medulla were identified and examined on a laser scanning confocal microscope (Zeiss 710 or 800). Cytoarchitectonic features of these nuclei and respective subnuclei were visualized and identified with the cytoplasmic neuronal marker HuC/D and used to define matching regions of interest (ROI) for quantitative analysis. Anatomically defined ROIs were used to quantify the density of labeled axons and terminals in the PVH and DVC. Additionally, cell type-specific targeting of AgRP inputs to PVH oxytocin neurons was analyzed by adapting a previously validated quantification protocol (4). In order to visualize and quantify AgRP inputs contacting PVH oxytocin neurons, sections were imaged at high magnification using an oil-immersion 63x objective, at a frequency of 0.0823 µm in the x and y planes and z-step of 0.427 µm. All other images were captured with a 20x objective, and collected at a frequency of 1.18 µm in the x and y planes, and a z-step of 1.38 µm, with 0.8x digital zoom unless otherwise noted.

Volocity visualization software was used to prepare 3D reconstructions of each multichannel set of images. To quantify the overall densities of labeled fibers in each ROI of the PVH and DVC, a previously validated methodology was utilized (4). Briefly, images were binarized and skeletonized such that fiber length is proportional to volume, and then total skeletonized fiber length was summed for each ROI throughout the volume of the image stack to estimate density of fibers within the ROI. The density of labeled inputs from AgRP neurons onto PVH oxytocin neurons, and density of Oxytocin-SynTom terminals in the DVC were quantified in this fashion. The terms ‘inputs ’or ‘terminals ’used in this study refer to the visualization of synaptophysin-tdTomato (SynTom) labeled axonal structures in close apposition to labeled neuronal cell bodies. The terms ‘fibers’,‘axons, ’or ‘projections ’include, but are not limited to SynTom-labeled axon structures, but also immunohistochemical labeling, which can be detected in axon terminals as well as fibers en passant. Quantification of these axonal elements were only considered to be in close apposition onto a target neuron in the PVH if they were touching by voxel-to-voxel contact, with no unlabeled voxels between the presynaptic and postsynaptic elements of the presumptive synapse. Super-resolution optical microscopy, or ultrastructural visualization, will be necessary to directly measure synaptic contacts onto postsynaptic cells, but the density of genetically targeted and immunohistochemically labeled inputs to oxytocin neurons appears to correlate with8 previous estimates of the density of synapses onto identified cell types within the PVH (4).

To measure the density of Fos and pSTAT3-immunoreactive nuclei in the ARH, PVH, and DVC induced by i.p. leptin injection, the number of Fos or pSTAT3-immunoreactive nuclei was counted in maximum projection confocal images through each region, guided by Volocity software.

### Experimental design and statistical analyses

Data are presented as group mean values ± SEM, as well as individual data points. Statistical analyses were performed using GraphPad Prism software (Version 9.5). Differences between groups were considered statistically significant at *p*<0.05. Significant differences between groups in response to cold challenge were tested with one-way repeated-measures ANOVA; unpaired t-tests were used for all other statistical tests. Sample sizes: AgRP-Cre-hM4Di + Sal: n = 7; AgRP-Cre-hM4Di + CNO: n = 7. Leptin-induced cFos assay: AgRP-Cre-hM4Di +Sal +Lep n = 3; AgRP-Cre-hM4Di +CNO +Lep n = 3. Cold tolerance test: Controls: n = 9; AgRP-Cre-hM4Di + CNO: n = 11. GI transit time and permeability assays: Control AgRP-Cre-hM4Di (no CNO) n = 5; AgRP-Cre-hM4Di +CNO: n = 6. Two-way repeated measures analysis of variance (ANOVA) with multiple comparisons was used to compare groups in the cold challenge assay. Student’s t-test was used to compare data within two groups, using paired and unpaired tests where appropriate.

Differences in AgRP density and oxytocin volume between genotypes was tested for significance by unpaired t-tests. One-way ANOVA followed by a pairwise post-hoc test was used to test for comparisons between three groups.

## Notes

### Competing Interest Statement

The authors have declared no competing interest.

